# Analysis of *Sp*Cas9 on- and off-target effects in high efficiency multiplex editing in Arabidopsis

**DOI:** 10.64898/2026.09.28.754964

**Authors:** Johannes Stuttmann, Jens Keilwagen, Philipe Ortet, Fang Shiang Lim, Thomas Schmutzer, Christel Llauro, Assane Mbodj, Marie Mirouze, Frank Hartung, Ralf Wilhelm

**Affiliations:** Institute for Biosafety in Plant Biotechnology, Julius Kühn-Institute - Federal Research Centre for Cultivated Plants (JKI), Erwin-Baur-Str. 27, 06484 Quedlinburg, Germany; Aix Marseille Univ, CEA, CNRS, BIAM, UMR7265, LEMiRE (Rhizosphère et interactions sol-plante-microbiote), 13115 Saint-Paul lez Durance, France; Institute of Agricultural and Nutritional Sciences, Martin Luther University Halle-Wittenberg, Karl-Freiherr-von-Fritsch-Straße 4, Halle (Saale) 06120, Germany; Laboratoire Génome et Développement des Plantes, UMR 5096 CNRS/UPVD, Université de Perpignan, Via Domitia, 52 Avenue Paul Alduy, 66860, Perpignan Cedex, France; IRD, EMR IRD-CNRS-UPVD ‘MANGO’, Université de Perpignan, Perpignan, France

**Keywords:** CRISPR/Cas, multiplexing, off-targets, whole genome sequencing, biotechnological safety

## Abstract

RNA-guided nucleases (RGNs), such as Cas9 from *Streptococcus pyogenes* (*Sp*Cas9), are widely used for plant genome editing. Previous surveys for off-targeting, the modification of unintended targets with similarity to the intended target, indicate high specificity of *Sp*Cas9 in plant cells. However, off-targeting has not been assessed for efficiency-optimized editing systems combined with extensive multiplexing, which might increase the likelihood of cleavage at unintended sites. We therefore analyzed *Arabidopsis thaliana* lines that had been extensively mutagenized using zCas9i and up to 29 gRNAs addressing >45 target sites over several rounds of editing. Genomes were sequenced by short- and long-read technologies, and genome-wide variants were catalogued. Our pipeline for variant calling reliably detected RGN-induced mutations at on-targets. When excluding these on-target modifications, variants were detected in edited lines at frequencies similar to those previously reported for spontaneous mutations. In further analyses, we did not find any evidence for an origin of these variants from RGN activity. Our data are thus consistent with high specificity of *Sp*Cas9. In contrast, we detected genomic reorganization events upon editing at two complex loci, *RPP1* and *RPP7*, encompassing multiple homologous genes, and also identified an allele by WGS that had escaped detection by amplicon sequencing. We conclude that, while off-targets may efficiently be avoided by selection of specific gRNAs, on-target modifications may be more extensive than intended, especially at complex loci and/or during multiplexing.

## Background

### Specificity of RNA-guided nucleases in genome editing applications

CRISPR/Cas system-derived RNA-guided nucleases (RGNs) are widely used in animal and plant genome editing. The specificity of RGNs is controlled by a variable region of the guide RNA (gRNA), commonly ∼20 nt in length, which engages in complementary base-pairing with the target DNA. Additionally, most RGNs require target sites (protospacers) to be flanked by a protospacer adjacent motif (PAM). For *Sp*Cas9 from *Streptococcus pyogenes*, the canonical PAM is NGG. To identify target sites, *Sp*Cas9 first interacts with low affinity with the PAM, after which the flanking sequence is scanned for complementarity to the gRNA, most likely involving some degree of lateral diffusion (Sternberg *et al*., 2014; Globyte *et al*., 2018). If sufficient complementarity exists, R-loop formation propagates directionally away from the PAM and the target sequence is cleaved.

The *Sp*Cas9 targeting mechanisms suggests high specificity, and full dependency of cleavage on presence of a PAM. Indeed, numerous studies in animal systems reported that off-targeting, the cleavage at loci with similarity to the true target, rarely occurs (Cho *et al*., 2014, Hsu *et al*., 2013, Pattanayak *et al*., 2013, Cho *et al*., 2013). In these analyses, off-targets were flanked by a PAM and had no more than three mismatches to the guide RNA. However, some studies also reported that off-targeting occurred at high frequencies, with some off-targets differing at up to six positions with the gRNA (Cradick *et al*., 2013, Fu *et al*., 2013).

In plants, multiple studies suggest high specificity of RGNs (Tang *et al*., 2018, Hahn & Nekrasov, 2019), and meta-analyses concluded that there was no evidence for off-targeting at sites with four or more mismatches (Modrzejewski *et al*., 2020; Sturme *et al*., 2022). However, previous analyses considered mainly sequences with similarity to the true target and flanked by a PAM sequence as potential off-targets, but cleavage of *Sp*Cas9 at several alternative PAMs was reported (Jiang *et al*., 2013; Zhang *et al*., 2014; Collias *et al*., 2020). Also, off-targeting was so far not analyzed in the context of efficiency-optimized editing systems and/or extensive multiplex editing, which might increase the likelihood for off-targeting by interfering effects.

We, therefore, decided to conduct a retrospective off-target analysis using lines of the model plant *Arabidopsis thaliana* (Arabidopsis) that were originally generated for a different purpose but, in the process, extensively mutagenized by multiplex editing and exposed to up to 29 gRNAs. We inspected both short- and long-read whole genome sequencing data to detect variants, and evaluated whether these might have resulted from off-targeting. Excluding on-target sites, we observed mutations (variants) at frequencies similar to those reported for spontaneous mutations in our edited lines (Ossowski *et al*., 2010; Monroe *et al*., 2022). Further analyses of variants did not support an origin from off-targeting. Accordingly, and in agreement with previous reports, our data are consistent with high specificity of *Sp*Cas9 even when efficiency-optimized systems and extensive multiplexing are combined. However, we observed genomic reorganization at on-target sites when editing complex loci containing multiple homologous genes with high similarity. Our analyses provide design considerations for future editing experiments, and we discuss our results in the light of the development of the implementing legislation for Category-1 genome edited plants in the EU.

## Results

### Plant Material

We previously generated a duodecuple (12x) mutant in Arabidopsis accession Col-0 using 24 gRNAs and efficiency-optimized zCas9i (Stuttmann *et al*., 2021; Grützner *et al*., 2021). We isolated a non-transgenic segregant (plant #1681-2) carrying bi-allelic mutations at all targeted loci (Figure 1a). Here, we sequenced two individuals descending from this plant, #1681-2-97 and #1681-2-60, by Illumina or Nanopore technologies, respectively (Figure 1a). Furthermore, a descendant from a sister plant that was not previously genotyped (#1681-3-99) was sequenced by both technologies (Figure 1a). Accordingly, the sequenced genomes of 1681-2/1681-3 descendants are separated by two generations.

**Figure 1:**
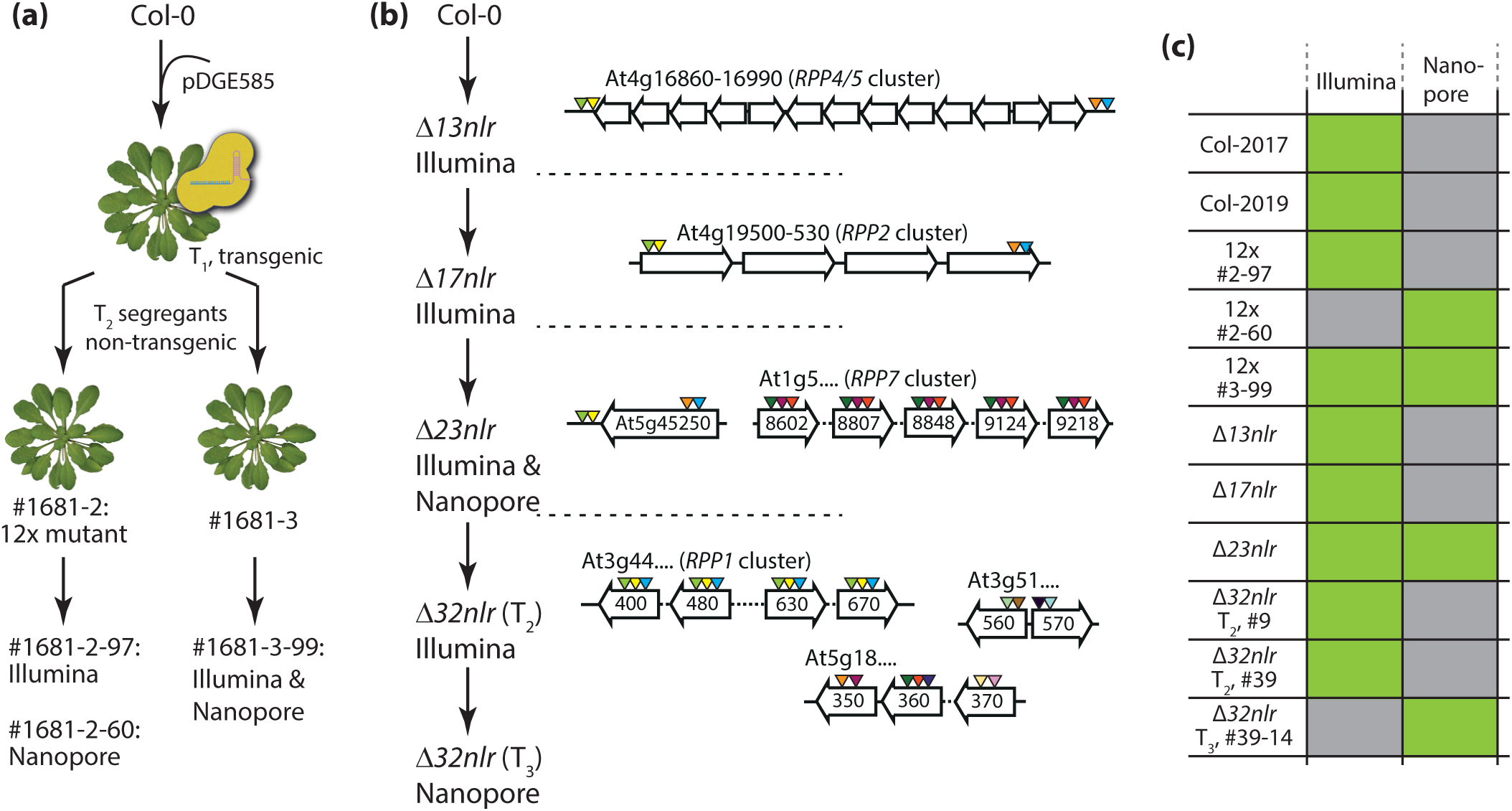
Plant material used for genome-wide off-target analyses. **a**) Genealogy of 12x mutant lines used for sequencing. Individual #1681-2 was previously characterized and carried bi-allelic mutations at all targeted loci. Two descendants were selected for sequencing by Illumina or Nanopore, respectively. A descendant from a sister plant that was not previously genotyped was sequenced by both Illumina and Nanopore technologies. **b**)D*nlr* mutant lines from consecutive editing rounds selected for sequencing. Each step consisted of at least three generations. Genes targeted in each step are shown; triangles indicate target sites (scheme not drawn to scale). **c**) Overview of plant material subjected to whole genome sequencing by short- and long-read technologies.

In a different set of experiments, we inactivated, by chromosomal deletion or point mutations, genes encoding NLR-(nucleotide binding – leucine-rich repeat)-type immune receptors (Figure 1b) (Van de Weyer *et al*., 2019; Bernoux *et al*., 2022). NLRs can recognize, directly or indirectly, so-called effector proteins translocated into plant cells by pathogenic microbes (Bernoux *et al*., 2022). Respective genes evolve rapidly in nature, and are commonly organized in clusters containing tandem arrays of highly similar genes (Van de Weyer *et al*., 2019). In iterative rounds of genome editing, we deleted the *RESISTANCE TO PERONOSPORA PARASITICA4/5* (*RPP4/5*) cluster (*Δ13nlr,* ∼ 80 kb) and the *RPP2* cluster (*Δ17nlr*; ∼ 30 kb) (Noël *et al*., 1999; Van Der Biezen *et al*., 2002; Sinapidou *et al*., 2004; Kim *et al*., 2014). Additional *NLR* genes including those at the complex *RPP7* and *RPP1* loci were targeted in two further rounds of editing to yield *Δ23nlr* and *Δ32nlr* lines (Figure 1b)(Chae *et al*., 2014; Stuttmann *et al*., 2016; Li *et al*., 2020; Ordon *et al*., 2021). In total, 29 gRNAs with more than 45 target sites - several gRNAs had multiple targets in homologous genes - were used (Figure 1b). Each consecutive round of editing consisted of at least three generations. Several individuals from different rounds of editing were selected for sequencing by Illumina and/or Nanopore technologies (Figure 1b, c).

We did not record a direct association of the CRISPR-generated mutant lines with Col-0 seed batches initially used for transformation. Therefore, we sequenced two Col-0 individuals derived from lab seed batches dated to 2017 (Col-2017) and 2019 (Col-2019; Figure 1c) as controls.

### Illumina sequencing and data analysis

The genomes of the nine selected lines were sequenced to moderate coverage (average 20x, min 17x, max 22x). After mapping, variant calling and initial filtering (Material and Methods), approximately 130 variants per genome remained and were analyzed in detail. We intended to treat any variant as a potential off-targeting result initially. We therefore called “off-target candidates” variants that were *i)* represented by at least four reads, *ii)* not supported by any reads in a genetically independent line, and *iii)* different from the TAIR10 reference (Tables 1, S1).

Upon comparing the genomes of the two wild type lines, Col-2019 and Col-2017, nine variants (two InDels, seven SNPs) were retained with these criteria (Table 1). The 12x mutants, as well as the *Δ13nlr* line, were compared against the two Col-0 genomes and TAIR10. After removal of on-targets, two, one and six InDels, and 17, 16 and 60 SNPs were retained as off-target candidates (Table 1). *Δ17nlr, Δ23nlr, Δ32nlr* were compared to *Δ13nlr.* Between 0-2 InDels and 4-7 SNPs were retained as off-target candidates for these lines (Table 1).

**Table 1:** Genome-wide detection of variants.

| | Col-2019 | 12x #2-97 | 12x #2-60 | 12x #3-99 | $\Delta 13\text{nlr}$<br>[#1203] | $\Delta 17\text{nlr}$<br>[#1253] | $\Delta 23\text{nlr}$<br>[#1767] | $\Delta 32\text{nlr}$<br>[#1933-9] | $\Delta 32\text{nlr}$<br>[#1943-39] | $\Delta 32\text{nlr}$<br>[#1943-39-14] |
| --- | --- | --- | --- | --- | --- | --- | --- | --- | --- | --- |
| InDels | 2 | 14 |  | 14 | 6 | 2 | 1 | 4 | 2 |  |
| InDels -<br>on-targets | n/a | 12 |  | 13 | 0 | 0 | 1 | 3 | 2 |  |
| InDels -<br>off-target candidates | **2** | **2** |  | **1** | **6** | **2** | **0** | **1** | **0** |  |
| SNPs -<br>off-target candidates | 7 | 17 |  | 16 | 60 | 6 | 4 | 7 | 7 |  |
| Insertions |  |  | 0 | 1 |  |  | 0 |  |  | 13 |
| Insertions -<br>on-targets |  |  | 0 | 0 |  |  | 0 |  |  | 13 |
| Insertions -<br>off-target candidates |  |  | **0** | **1** |  |  | **0** |  |  | **0** |
| Deletions |  |  | 7 | 9 |  |  | 7 |  |  | 17 |
| Deletions -<br>on-targets |  |  | 5 | 7 |  |  | 3 |  |  | 14 |
| Deletions -<br>off-target candidates |  |  | **2** | **2** |  |  | **4** |  |  | **3** |
| Duplications |  |  | 0 | 0 |  |  | 0 |  |  | 1 |
| Duplications -<br>on-targets |  |  | 0 | 0 |  |  | 0 |  |  | 1 |
| Duplications -<br>off-target candidates |  |  | **0** | **0** |  |  | **0** |  |  | **0** |
| Inversions |  |  | 0 | 0 |  |  | 0 |  |  | 1 |
| Inversions -<br>on-targets |  |  | 0 | 0 |  |  | 0 |  |  | 1 |
| Inversions -<br>off-target candidates |  |  | **0** | **0** |  |  | **0** |  |  | **0** |

Overall, similar numbers of SNPs and InDels were thus detected upon comparison of the two Col-0 lines and in CRISPR lines. The *Δ13nlr* line was a notable exception (Table 1). Intriguingly, most of the variants detected in this line (64/66) were homozygous, which would not be expected for newly induced variants. We therefore considered that *Δ13nlr* had derived from an inbred Col-0 lineage different from the sequenced control lines. We tested this hypothesis by analyzing polymorphisms detected in *Δ13nlr* in individual plants grown from further available Col-0 seed batches. Indeed, we identified a plant that carried the same haplotype as *Δ13nlr* at 10/13 tested loci, distributed over all five chromosomes (Appendix 1). This strongly supports that indeed a different inbred Col-0 population was used as starting material for induction of *Δnlr* mutations.

### Nanopore sequencing and data analysis

Four genomes (Figure 1c) were sequenced on individual MinION Flow Cells, resulting in an average coverage of 43x (min 23x, max 70x) per genome. We used Sniffles2 (Smolka *et al*., 2024) for detection of structural variants of a minimal length of ten nucleotides (nt; Table S2). Datasets were intersected and filtered, and variants were further inspected using IGV. 7-32 variants per genome were retained, 80% (45/56) of which corresponded to mutations at on-target sites (Table S3). Eleven variants, all of which were deletions, remained as off-target candidates.

### Benchmarking of variant calling parameters

We used the detection of mutations at on-targets to benchmark our variant detection workflow. Individual #1681-2, from which #2-97 and #2-60 were derived, was previously genotyped by amplicon sequencing (Stuttmann *et al*., 2021). We used short- and long-read data to reconstruct the genotype of the parental line, and observed inconsistencies at two loci (Table 2). In one case (At4g26090), a haplotype demonstrating -21 and -102 nt deletions was apparently missed previously. Further, a complex allele resulting in deletion of ∼ 750 nt at At3g27920 could not be detected by amplicon sequencing, as it resulted in deletion of the primer binding site (Figure S1). We also deduced the genotype of individual #3-99 (Table 2). All on-target mutations were InDels, as expected (Vu *et al*., 2017; Shen *et al*., 2017).

**Table 2:**
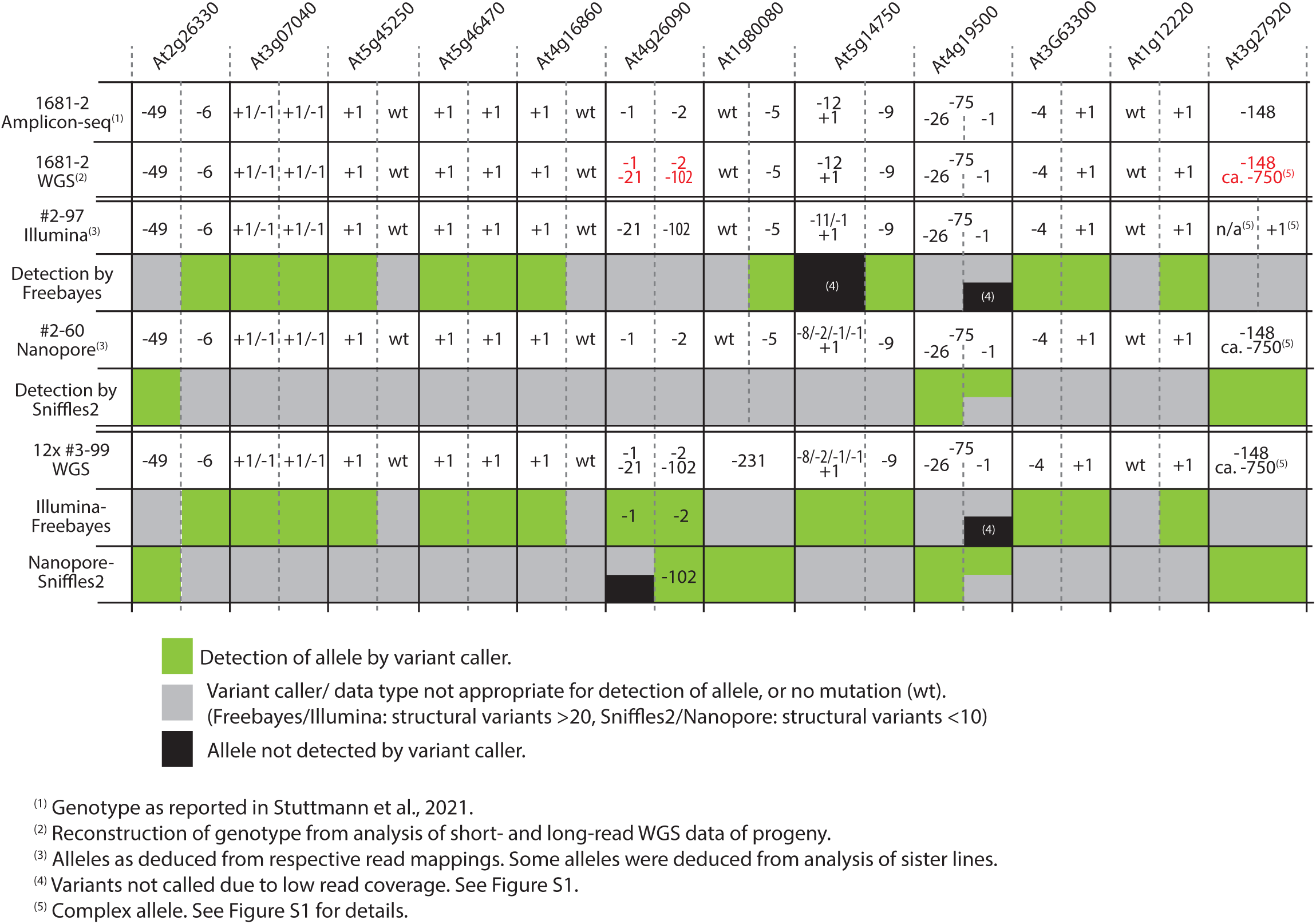
Genotype(s) and variant calling in 12 x mutant lines.

With the parameters we used for variant calling, InDels ≤20 nt could be detected in short read data (Illumina, Freebayes), and ≥10 nt in long read data (Nanopore, Sniffles2). Overall, 38/43 of the variants at on-targets were detected by at least one variant caller (Table 2). For the remaining variants, in 4/5 cases, local perturbation of the read mappings due to neighboring/overlapping on-target mutations from multiplexing impeded detection (Figure S2). The occurrence of several mutations in close proximity from off-targeting is highly unlikely. Accordingly, we conclude that our variant detection workflow can detect off-targeting with high fidelity.

### Off-target analysis

We wanted to evaluate whether the detected variants (“off-target candidates”, Table 1) could result from off-targeting, or rather represented spontaneous mutations. We extracted variants and surrounding sequence (50 nt up- and downstream; Tables S4, S5), and searched for homology between these genomic regions and the gRNAs used for editing in respective lines. We used the Levenshtein distance (Levenshtein, 1966) to identify gRNAs most closely related to any sequence stretch within the extracted genomic regions. The Levenshtein distance refers to the minimal number of transformations (exchanges, insertions, deletions) that is required to convert one string of symbols into another. Accordingly, we searched for the gRNA that could be converted into a sequence within the extracted genomic regions with the minimal number of transformations.

On average, more than six transformations were required to convert gRNAs into a sequence within the extracted regions (121 genomic regions representing ∼32 kb were included in analyses). We inspected InDels more closely, as this is the variant type predominantly induced by RGN mutagenesis (Table 2) (Lemos *et al*., 2018; Chen *et al*., 2019). Across all experiments, 4-7 transformations were required to convert a gRNA into a sequence within one of the extracted regions (Figure 2, Table S5). None of these sequences was flanked by a canonical or non-canonical PAM. Our data do thus not support an origin of these variants from RGN mutagenesis and off-targeting.

**Figure 2:**
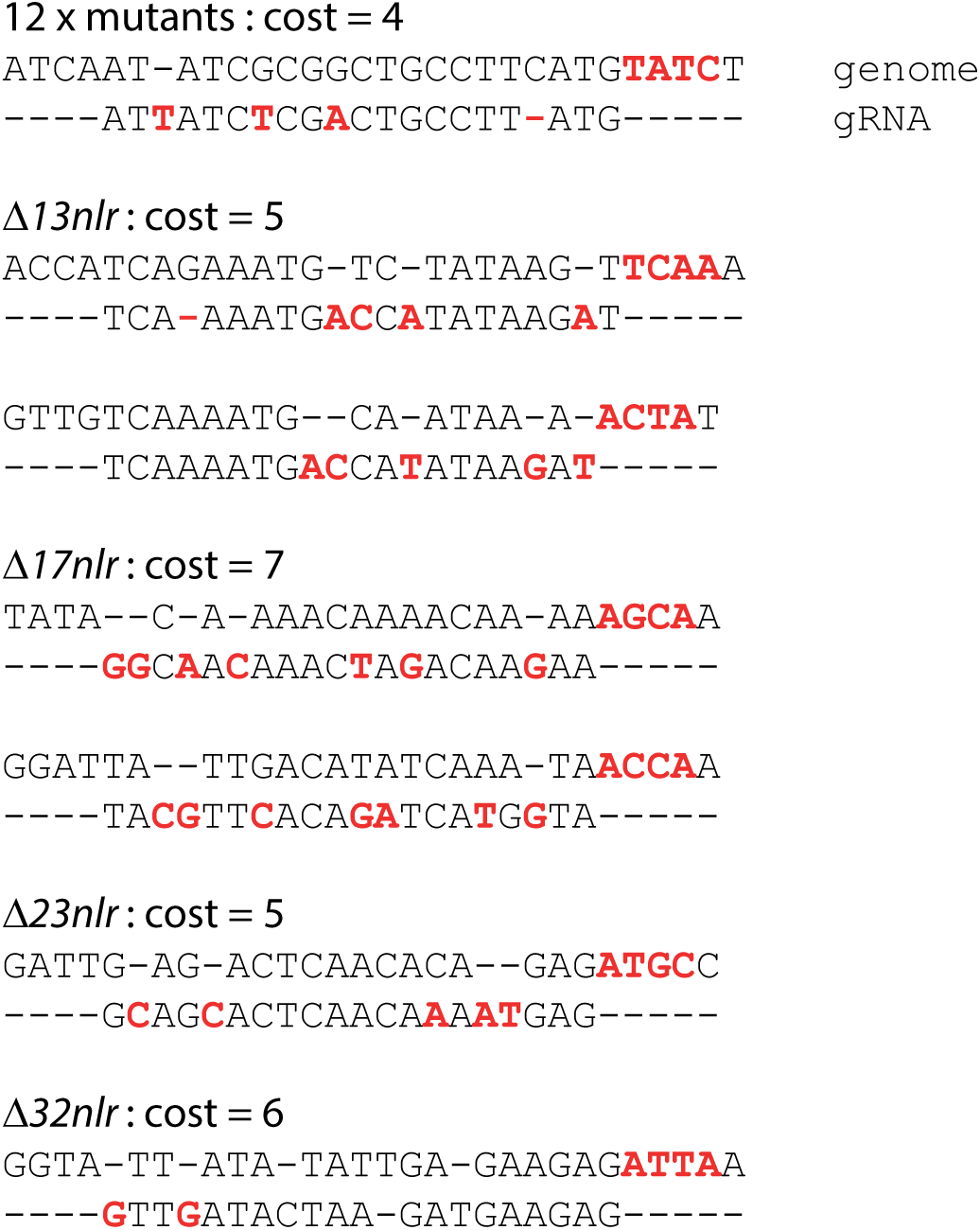
Best matches between gRNAs and variant-containing regions as determined by Levenshtein distance. Genomic regions containing variants were extracted (Tables S4, S5), and compared to gRNA sequences of respective editing experiments using the Levenshtein distance. The best matches between regions containing InDels and gRNAs are shown for the different editing experiments, as indicated. Transformations that are required to convert the gRNA into the genomic sequence are marked in red on the gRNAs. The position at which a PAM sequence would be expected for cleavage by *Sp*Cas9 is marked in red on the genomic sequence. The „best match(es)“ is/are shown, „cost” indicates the number of required transformations.

We also scrutinized off-targets predicted by CRISPOR (Table S6)(Concordet & Haeussler, 2018). Predicted off-targets did not coincide with any of the detected variants. Further, when inspecting read mappings in IGV, no variants were observed at any of the predicted off-targets. Summarizing, we do not find any evidence for off-targeting in our sequencing data.

### Genomic rearrangements at the *RPP7* and *RPP1* on-target loci

The *RPP7* and *RPP1* loci are complex, comprising four and five highly similar NLR-coding genes, respectively. In our experiments, each locus was targeted with three gRNAs at conserved target sites within the NLR-coding genes, resulting in a total of 15 and 12 target sites, respectively (Figure 1b). Inspection of these loci in IGV suggested extensive mutagenesis in *Δnlr* lines.

At the *RPP7* locus, no variants were called, but read mappings showed multiple break points and suggested several larger deletions. However, due to highly repetitive sequences at this locus, read mappings were of low quality (Figure S3a). Manual analysis of Nanopore read ends suggested an inversion, fusing At1g58602 with the 5’ section of either At1g58807 or At1g59124 (Figure S3b). We attempted to gain further insights into recombination events by *de novo* assembly of Nanopore reads from this region, and comparison of the *de novo* assembly to TAIR10 (Figure S3c, d). For a control line, this led to a minor expansion of the region, but no major discrepancies (Figure S3c). By contrast, for the *Δ32nlr* line, the same analysis suggested multiple translocations, inversions and deletions (Figure S3d), and the assembled region lacked approximately 12 kb in comparison to the control. We did not further attempt to resolve the haplotype present at *RPP7* in *Δ32nlr* due to its complexity. However, we conclude that our CRISPR mutagenesis involving many target sites in a region containing the highly similar RPP7 genes likely resulted in genomic reorganization.

At the *RPP1* locus, in total 21 variants, including insertions, deletions, inversions and a large duplication were called in *Δ32nlr* by Sniffles (Table S3). Read coverage along the *RPP1* region did not support a large duplication, and mappings showed multiple breakpoints and a possible deletion (Figure 3a). As for *RPP7*, we extracted Nanopore reads and conducted *de novo* assembly. For the control, this resulted in three contigs, which showed overall high congruency with the TAIR10 reference (Figure 3b). In contrast, for *Δ32nlr*, the *de novo* assembly suggested a deletion of ∼ 45 kb, and two inverted regions of ∼ 100 and ∼ 20 kb (Figure 3c). We designed primers for presence-absence analyses and amplicons across breakpoints to further decipher the haplotype of *Δ32nlr* at the *RPP1* locus (Figure 3a). Initial analyses at At3g44630 suggested that the individual subjected to Nanopore sequencing was heterozygous for two different alleles (Figure 3d), and we indeed observed segregation when individuals from the following generation were analyzed (Figure S5). Two homozygous individuals were selected for further analyses, alongside controls (Figure S5).

**Figure 3:**
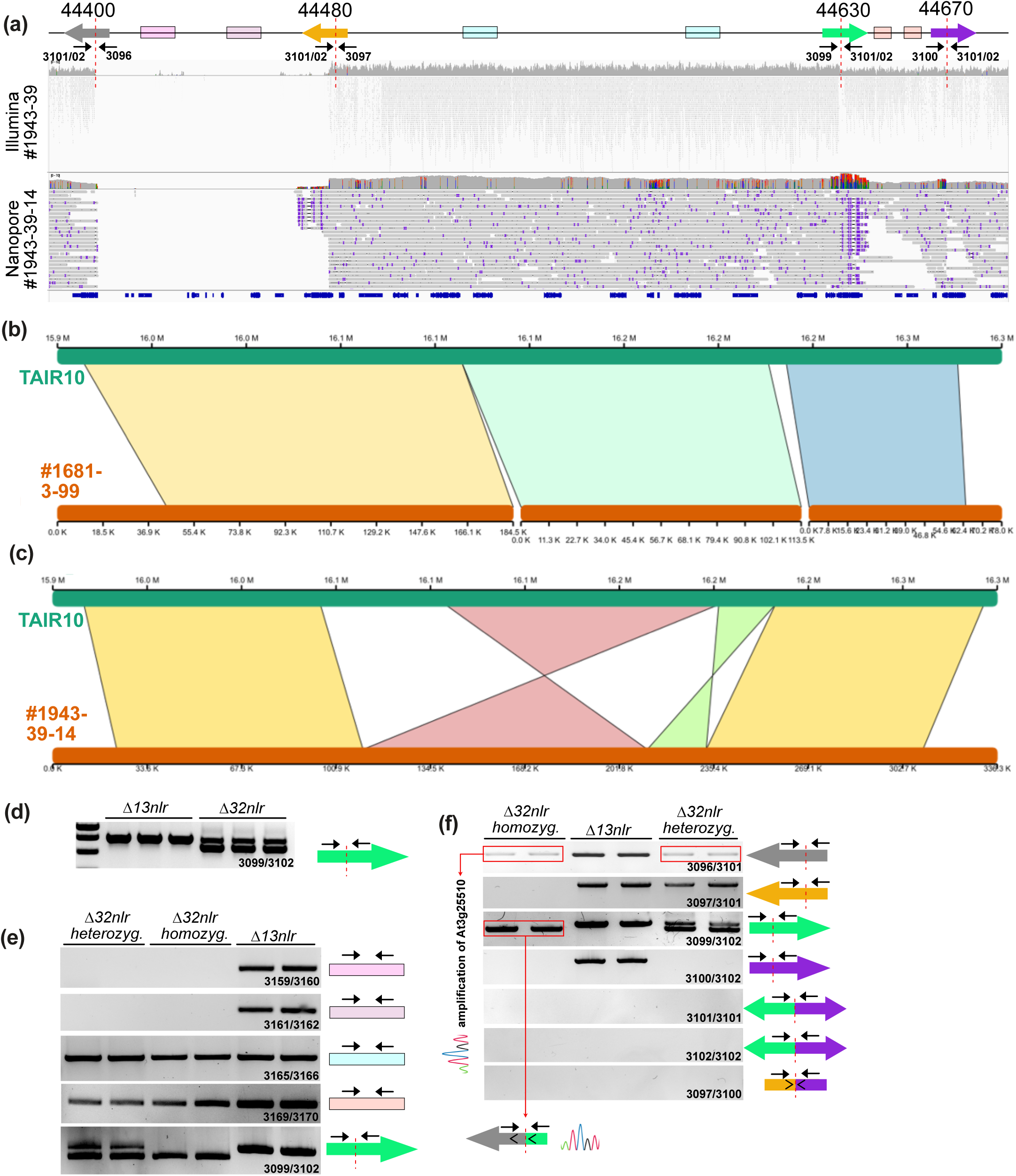
Analysis of recombination events at the *RPP1* locus. **a**) Mappings of Illumina and Nanopore reads from plants #1943-39 and #1943-39-14 visualized in IGV. The scheme on top illustrates primer binding sites as well as regions queried by PCR genotyping. **b**) Comparison of a *de novo* alignment of Nanopore reads from the *RPP1* region for a control line, #1681-3-99, which was not edited at the *RPP1* locus. **c**) As in b), but reads from the Δ*32nlr* line #1943-39-14 were used for the *de novo* assembly. **d**) PCR-analysis of the breakpoint within At3g44400, suggesting that the Nanopore-sequenced Δ*32nlr* individual #1943-39-14 bears two different haplotypes at the *RPP1* locus. **e**) Presence-absence PCRs on regions between breakpoints within the *RPP1* locus in Δ*32nlr*. **f**)PCR-analysis of breakpoints within RPP1-like genes in Δ*32nlr*. The weak amplicon for oligonucleotides 3096/3101 in Δ*32nlr* individuals was identified as belonging to At3g25510 by Sanger sequencing. The same amplicon from the control line was derived from At3g44400, as expected. Sequencing of the amplicon obtained with oligonucleotides 3099/3102 from Δ*32nlr* individuals homozygous at the *RPP1* locus revealed a inversion and fusion event involving At3g44400 and At3g44630; details are provided in Figure S5.

Presence-absence analyses supported a deletion of the segment in between At3g44400 and At3g44480, as suggested by visualization of read mappings and also analysis of the *de novo* assembly (Figure 3a, c, e). Similarly in agreement with the *de novo* assembly, we obtained evidence for an inversion and fusion involving At3g44400 and At3g44630 (Figures 3f, S6). However, using different primer combinations, we could not resolve additional junctions. Accordingly, we could not resolve the haplotype that was induced by editing at the *RPP1* locus, but our data suggest substantial genomic reorganization.

### Discussion and Outlook

In June 2026, the European Parliament and the Council approved Regulation 2026/1388 about genome-edited plants, considerably reducing the regulatory burden compared to transgenic plants (http://data.europa.eu/eli/reg/2026/1388/oj). The regulation defines two categories of NGT plants: Category-1 plants are deemed equivalent to conventionally bred varieties and pass a formal verification before release into the environment. Category-2 plants still have to be authorized, undergo a more adjusted risk assessment process, and further obligations remain similar to those applied to transgenic plants. Especially for Category-1 NGT plants, this is a landmark shift after the ECJ ruling had subjected all gene-edited plants to the strict GMO regulations in 2018. During a two-year implementation period, the European Commission will be developing the necessary legislation and guidance for implementation. Market releases under the new regulation can be expected from mid-2028. We discuss our analyses of highly mutagenized Arabidopsis lines in the light of the development of this implementing legislation.

### Off-targets – and avoiding them

Off-target effects will stay a subject of discussion with regard to Category-1 crop development. The legislation defines a maximum of 20 modifications for Category-1 plants – it is currently not known whether these shall include possible off-target mutations, how these may be defined and which verification criteria will need to be fulfilled.

In the off-target survey conducted here, we retained all variants detected in edited lines as off-target candidates to avoid applying any biases (Table 1). However, there is no evidence for non-sequence-specific cleavage of DNA by RGNs under *in vivo* conditions (Sundaresan *et al*., 2017). Accordingly, potential off-targets must share significant sequence similarity with the intended target, which allows their cataloguing and evaluation *a priori*, provided that genome information for the cultivar of interest is available. The CFD and MIT scoring systems are most commonly used to evaluate off-targets (Hsu *et al*., 2013; Doench *et al*., 2016). Sites with ≤ 4 mismatches are generally taken into account and further restrictions resulting from the employed nuclease may be applied, such as, *e.g.*, retaining only sites flanked by NGG, NAG or NGA when *Sp*Cas9 is used. A previous analysis concluded that a minimal CFD score of 0.023 removes 57% of false positives while losing only 2% of true positives, and that at this cut-off, no off-target with a modification frequency above 1% was missed (Haeussler *et al*., 2016). However, such a sensitivity-maximizing cut-off value would not be reasonable for the collection of possible off-targets in plant genome editing applications. First, the mentioned conclusions were derived from animal cell experiments and a training set biased toward promiscuous guides (Haeussler *et al*., 2016). Further, available off-target surveys within the plant kingdom suggest high specificity of RGNs in plant cells: Meta-analyses previously conducted by us and others concluded that there is no relevant evidence for off-targeting at sites with four or more mismatches (Modrzejewski *et al*., 2020; Sturme *et al*., 2022). In agreement, more than 100 predicted off-targets in editing experiments conducted here had CFD scores >0.2 (up to 0.93), and mutations were not observed at these sites (Table S6). In one implementation, all sites with ≤3 mismatches to a gRNA could be considered as off-targets that require verification. However, a mere mismatch count would, on the one hand, over-estimate potential off-targets, and, on the other hand, miss true off-targets resulting, *e.g.*, from rG:dT (gRNA:target DNA) pairings that do not provoke any strong penalty. Therefore, a better solution would be the definition of a cut-off score for, *e.g.*, the CFD score, to define off-target candidates that require verification in an edited line. Such a value could be derived from the totality of off-targeting events confirmed in plant genome editing to date, by identifying the cut-off that retains the large majority of these events while minimizing the number of candidate sites requiring verification. Given the limited number of confirmed events, such a cut-off would initially be provisional and should be subject to review as further plant data accumulate. In either case, when genome information is available, researchers and breeders are able to select high-specificity gRNAs with ease using bioinformatic tools to minimize the risk of inducing undesired mutations.

The situation becomes more complicated when genome information is not available. In this case, unbiased *in vitro* approaches such as CIRCLE-seq (Kim *et al*., 2015; Cameron *et al*., 2017; Tsai *et al*., 2017), with a standardized protocol, could be employed to query gRNA specificity and catalogue potential off-target sites prior to *in vivo* editing. However, it should be noted that complexity and cost of such analyses scale with genome size, and that many important crops, especially polyploid cereals, have very large genomes.

### Verification of on-targets

We detected genomic reorganization events at two complex loci encompassing multiple homologous genes – the *RPP1* and *RPP7* loci encoding NLRs (Figures 3, S4). Further, we noted that standard PCR genotyping may miss on-target alleles (here at the At3g27920 locus; Table 2, Figure S1) due to deletion of the primer binding site. Notably, respective *NLR* loci were previously identified as recombination hotspots (Jiao & Schneeberger, 2020). In animal cells, dedicated experiments were employed to detect on-target mutations that are more complex than insertion/deletion of a few nucleotides – mostly large deletions (Kosicki *et al*., 2018, 2022; Geng *et al*., 2022; Park *et al*., 2022). In plants, large deletions were induced intentionally in several cases including the present study (Zhou *et al*., 2014; Ordon *et al*., 2017; Duan *et al*., 2021; Ordon *et al*., 2023). Also, multiplex editing, especially at tandemly arrayed genes, has been previously associated with the occurrence of deletion-inversion bi-alleles that may complicate genotyping (Liu *et al*., 2023). However, to the best of our knowledge, there is little evidence for complex on-target modifications including large deletions at single target sites, and these have not been systematically studied in plants (Sturme *et al*., 2022). Nonetheless, our observations emphasize that on-target verification by sequencing of short PCR amplicons will not in all cases be sufficient to determine the precise haplotype(s) induced by RGN mutagenesis.

The simplest solution could be the definition of a mandatory “verification window” ensuring the detection of the majority of RGN-induced on-target mutations. However, data providing a rationale for the definition of such a window cannot be derived from animal cell experiments and are scarce for the plant kingdom. Long-read sequencing represents an efficient and unbiased genotyping method, and could be coupled with target enrichment strategies for, *e.g.*, large cereal genomes. Alternatively, a multi-stage approach could be defined to verify on-target mutations. At first instance, considering non-chimeric segregants, sequencing may identify a number of alleles that unambiguously establishes the induced haplotype. To that end, *e.g.*, detection of two different alleles in a diploid species evidences that all possible alleles were detected. If a single allele is detected in the same scenario, the individual could either be homozygous for the allele, or a second allele might have been missed. Quantitative PCR techniques comparing the abundance of the detected allele with control loci can discriminate between these different cases. The interrogated region can subsequently be extended if quantitative PCR suggests that not all alleles were detected.

Base editing and prime editing carry their own risks, such as target-independent deamination in the case of base editors, and the induction of DNA nicks. However, neither technique induces a targeted double-strand break as the intended outcome, and both may thus be preferable for the modification of complex loci encompassing repetitive sequences prone to recombination, at which the genomic reorganizations reported here were observed. It may therefore be reasonable to define different verification requirements according to the distinct risks associated with the employed editing modalities.

## Material and Methods

### Plant material, growth conditions and transformation

*Arabidopsis thaliana* accession Columbia-0 (Col-0) plants were cultivated under short day conditions in a walk-in chamber (8h light, 23/21°C day/night, 60 % relative humidity) or in a greenhouse under long day conditions (16h light) for seed set. Arabidopsis was transformed by floral dipping as previously described (Logemann *et al*., 2006). Agrobacterium strain GV3101 pMP90 was used. Primary transformants (T_1_) were selected by seed fluorescence (Shimada *et al*., 2010) using a stereomicroscope equipped with an mRFP filter. In the T_2_ generation, non-fluorescent seeds were selected, respective plants genotyped and propagated to the next generation.

### Guide RNA design and molecular cloning

Target sites were selected using CRISPOR and/or ChopChop (Labun *et al*., 2016; Concordet & Haeussler, 2018); gRNAs are provided in Tables S4 and S6. gRNAs and final plant transformation vectors were assembled as previously described (Ordon *et al*., 2017, 2021).

### Illumina sequencing and data analysis

DNA for Illumina sequencing was extracted from single plants using the Qiagen DNeasy Plant Kit according to manufacturer’s instructions. DNA was quantified on a Qubit, and sequenced by Genewiz (Azenta Life Sciences); 2 x 150 nt paired-end reads. Raw data was trimmed with Trim Galore (https://github.com/FelixKrueger/TrimGalore) using non-default parameters --quality 30 and --length 50. Data was mapped against the *Arabidopsis thaliana* TAIR10 reference genome using bowtie2 v2.4.2 (Langmead & Salzberg, 2012). Mapped reads were filtered with samtools (version 1.10) excluding unmapped, not primary or supplementary alignments. Further, duplicate reads were removed. Raw variant calling was done with FreeBayes v1.3.1-dirty (Garrison & Mart, 2012). A custom script was used to filter raw variant calls. Variants with quality of less than 40 were removed. In addition, variant calls of a sample with coverage of less than 6 reads or more than 100 were set to unknown.

### Nanopore sequencing and data analysis

DNA was extracted following the “arabidopsis-leaf-dna” protocol from Nanopore (<u>link</u>) from 0,6 g of starting material with Carlson lysis buffer (100 mM Tris-HCl, pH 9.5, 2% CTAB, 1.4 M NaCl, 1% PEG 8000, 20 mM EDTA) and using the Qiagen Blood & Cell Culture DNA Midi Kit for purification. Size selection with SSB2X buffer was conducted to enrich large fragments, and size-selected ssDNA was quality-checked by agarose gel electrophoresis. Libraries were prepared with ONT Kit (SQK-LSK109) according to manufacturer’s instructions. Libraries were sequenced in house (UPVD Bio-Environment sequencing platform) on ONT MinION using R9.4.1 flow cells. Base-calling was done using Guppy (Oxford Nanopore Technologies) with high-accuracy mode, and assembled to the TAIR10 reference using minimap (Li, 2018) with default parameters. Read alignments were inspected for the presence of structural variations using Sniffles2 (Smolka *et al*., 2024). Default parameters were used, except that minimal length to integrate insertions and deletion (INDELs) was set ≥ 10. On average, ∼ 2900 variants per genome were obtained. We then intersected variants with those of genetically independent lines: all variants with coordinates that were identical of differed by ≤ 20 nt were excluded (Figure S1). Further, variants in centromeric regions (Rabanal *et al*., 2022) were excluded, and also those with allele frequency <0.3. Remaining variants (∼70 per genome) were inspected in read mappings using IGV.

### Resolving structural rearrangements by targeted read extraction and de novo assembly

Nanopore reads were extracted from mappings between the coordinates Chr1:21,720,000– 21,870,000 and Chr3:16,000,000–16,250,000 corresponding, respectively, to the *RPP7* and *RPP1* loci and approximately 20kb flanking sequence for the *Δ32nlr* line and a control line, #1681-3-99. In total, 297 and 652 reads were extracted from the *rpp7* and *rpp1* loci for *Δ32nlr*. Reads were subjected to *de novo* assembly using Flye v2.8.2 (Kolmogorov *et al*., 2019). Assembled contigs were realigned to the TAIR10 reference genome using NUCmer v4.0.0 (Marçais *et al*., 2018) from the MUMmer package, with the --maxmatch option and a minimum alignment length -L of 2000 bp, as recommended for highly repetitive regions. NGenomeSyn v1.40 (He *et al*., 2023) was used to visualize synteny between the assembled contigs and the reference genome.

## LIST OF MATERIAL

Figure 1: Plant material used for genome-wide off-target analyses.

Figure 2: Best matches between gRNAs and variant-containing regions as determined by Levenshtein distance.

Figure 3: Analysis of recombination events at the *RPP1* locus.

Supplemental Figure S1: Read mappings at the *GL1* locus for 12x mutant lines.

Supplemental Figure S2: Read mappings at the *RPP2* and *WER1* loci for 12x mutant lines.

Supplemental Figure S3: Analysis of recombination events at the *RPP7* locus.

Supplemental Figure S4: PCR-screening to identify plants homozygous for a single haplotype at the *RPP1* locus.

Supplemental Figure S5: Inversion and fusion event involving At3g44630 and At3g44400 in *Δ32nlr*.

Supplemental Figure S6: Intersection of variants detected by Sniffles2.

Table S1: Variants called on Illumina data.

Table S2: All variants called on Nanopore data.

Table S3: Final variants retained from analysis of Nanopore data.

Table S4: Compilation of all “off-target candidates” and gRNAs used in this study.

Table S5: Sequences extracted and used for Levenshtein distance analysis.

Table S6: Off-targets predicted by CRISPOR.

Appendix 1 Analysis of polymorphisms in a Col-0 individual

## Data Availability Statement

Sequencing data associated with this study was deposited in the European Nucleotide Archive under project accession PRJEB123131.

## Author contributions

JS, RW, FH and JK designed the study. JS performed experiments and analyzed data. CL, AM and MM conducted Nanopore sequencing and analyzed data. TS provided additional analyses, and PO assisted in intersection and filtering. JK and FSL conducted remaining bioinformatic analyses. JS analyzed data, prepared figures and wrote the paper, all authors contributed to the final version.

## Ethical Approval

Not applicable.

## References

Bernoux M, Zetzsche H, Stuttmann J. 2022. Connecting the dots between cell surface- and intracellular-triggered immune pathways in plants. Current Opinion in Plant Biology 69: 102276.

Cameron P, Fuller CK, Donohoue PD, Jones BN, Thompson MS, Carter MM, Gradia S, Vidal B, Garner E, Slorach EM, et al. 2017. Mapping the genomic landscape of CRISPR–Cas9 cleavage. Nature Methods 14: 600–606.

Chae E, Bomblies K, Kim S-T, Karelina D, Zaidem M, Ossowski S, Martín-Pizarro C, Laitinen RAE, Rowan BA, Tenenboim H, et al. 2014. Species-wide Genetic Incompatibility Analysis Identifies Immune Genes as Hot Spots of Deleterious Epistasis. Cell 159: 1341–1351.

Chen W, McKenna A, Schreiber J, Haeussler M, Yin Y, Agarwal V, Noble WS, Shendure J. 2019. Massively parallel profiling and predictive modeling of the outcomes of CRISPR/Cas9-mediated double-strand break repair. Nucleic Acids Research 47: 7989–8003.

Collias D, Leenay RT, Slotkowski RA, Zuo Z, Collins SP, McGirr BA, Liu J, Beisel CL. 2020. A positive, growth-based PAM screen identifies noncanonical motifs recognized by the S. pyogenes Cas9. Science Advances 6: eabb4054.

Concordet J-P, Haeussler M. 2018. CRISPOR: intuitive guide selection for CRISPR/Cas9 genome editing experiments and screens. Nucleic Acids Research 46: W242–W245.

Doench JG, Fusi N, Sullender M, Hegde M, Vaimberg EW, Donovan KF, Smith I, Tothova Z, Wilen C, Orchard R, et al. 2016. Optimized sgRNA design to maximize activity and minimize off-target effects of CRISPR-Cas9. Nature Biotechnology 34: 184–191.

Duan K, Cheng Y, Ji J, Wang C, Wei Y, Wang Y. 2021. Large chromosomal segment deletions by CRISPR/LbCpf1-mediated multiplex gene editing in soybean. Journal of Integrative Plant Biology 63: 1620–1631.

Garrison E, Mart G. 2012. Haplotype-based variant detection from short-read sequencing.

Geng K, Merino LG, Wedemann L, Martens A, Sobota M, Sanchez YP, Søndergaard JN, White RJ, Kutter C. 2022. Target-enriched nanopore sequencing and de novo assembly reveals co-occurrences of complex on-target genomic rearrangements induced by CRISPR-Cas9 in human cells. Genome Research 32: 1876–1891.

Globyte V, Lee SH, Bae T, Kim J, Joo C. 2018. CRISPR/Cas9 searches for a protospacer adjacent motif by lateral diffusion. The EMBO Journal 38: EMBJ201899466.

Grützner R, Martin P, Horn C, Mortensen S, Cram EJ, Lee-Parsons CWT, Stuttmann J, Marillonnet S. 2021. High-efficiency genome editing in plants mediated by a Cas9 gene containing multiple introns. Plant Communications 2: 100135.

Haeussler M, Schönig K, Eckert H, Eschstruth A, Mianné J, Renaud J-B, Schneider-Maunoury S, Shkumatava A, Teboul L, Kent J, et al. 2016. Evaluation of off-target and on-target scoring algorithms and integration into the guide RNA selection tool CRISPOR. Genome Biology 17: 148.

He W, Yang J, Jing Y, Xu L, Yu K, Fang X. 2023. NGenomeSyn: an easy-to-use and flexible tool for publication-ready visualization of syntenic relationships across multiple genomes. Bioinformatics 39: btad121.

Hsu PD, Scott DA, Weinstein JA, Ran FA, Konermann S, Agarwala V, Li Y, Fine EJ, Wu X, Shalem O, et al. 2013. DNA targeting specificity of RNA-guided Cas9 nucleases. Nature Biotechnology 31: 827–832.

Jiang W, Bikard D, Cox D, Zhang F, Marraffini LA. 2013. RNA-guided editing of bacterial genomes using CRISPR-Cas systems. Nature Biotechnology 31: 233–239.

Jiao W-B, Schneeberger K. 2020. Chromosome-level assemblies of multiple Arabidopsis genomes reveal hotspots of rearrangements with altered evolutionary dynamics. Nature Communications 11: 989.

Kim D, Bae S, Park J, Kim E, Kim S, Yu HR, Hwang J, Kim J-I, Kim J-S. 2015. Digenome-seq: genome-wide profiling of CRISPR-Cas9 off-target effects in human cells. Nature Methods 12: 237–243.

Kim DS, Li Y, Ahn H-K, Woods-Tör A, Cevik V, Furzer OJ, Ma W, Tör M, Jones JDG. 2014. ATR2Cala2 from Arabidopsis-infecting downy mildew requires 4 TIR-NLR immune receptors for full recognition.

Kolmogorov M, Yuan J, Lin Y, Pevzner PA. 2019. Assembly of long, error-prone reads using repeat graphs. Nature Biotechnology 37: 540–546.

Kosicki M, Allen F, Steward F, Tomberg K, Pan Y, Bradley A. 2022. Cas9-induced large deletions and small indels are controlled in a convergent fashion. Nature Communications 13: 3422.

Kosicki M, Tomberg K, Bradley A. 2018. Repair of double-strand breaks induced by CRISPR–Cas9 leads to large deletions and complex rearrangements. Nature Biotechnology 36: 765–771.

Labun K, Montague TG, Gagnon JA, Thyme SB, Valen E. 2016. CHOPCHOP v2: a web tool for the next generation of CRISPR genome engineering. Nucleic Acids Research 44: W272–W276.

Langmead B, Salzberg SL. 2012. Fast gapped-read alignment with Bowtie 2. Nature methods 9: 357– 359.

Lemos BR, Kaplan AC, Bae JE, Ferrazzoli AE, Kuo J, Anand RP, Waterman DP, Haber JE. 2018. CRISPR/Cas9 cleavages in budding yeast reveal templated insertions and strand-specific insertion/deletion profiles. Proceedings of the National Academy of Sciences 115: E2040–E2047.

Levenshtein VI. 1966. Binary Codes Capable of Correcting Deletions, Insertions, and Reversals. 10.

Li H. 2018. Minimap2: pairwise alignment for nucleotide sequences. Bioinformatics 34: 3094–3100.

Li L, Habring A, Wang K, Weigel D. 2020. Atypical Resistance Protein RPW8/HR Triggers Oligomerization of the NLR Immune Receptor RPP7 and Autoimmunity. Cell Host & Microbe 27: 405–417.e6.

Liu J, Wang F-Z, Li C, Li Y, Li J-F. 2023. Hidden prevalence of deletion-inversion bi-alleles in CRISPR-mediated deletions of tandemly arrayed genes in plants. Nature Communications 14: 6787.

Logemann E, Birkenbihl RP, Ülker B, Somssich IE. 2006. An improved method for preparing Agrobacterium cells that simplifies the Arabidopsis transformation protocol. Plant Methods 2: 16.

Marçais G, Delcher AL, Phillippy AM, Coston R, Salzberg SL, Zimin A. 2018. MUMmer4: A fast and versatile genome alignment system. PLOS Computational Biology 14: e1005944.

Modrzejewski D, Hartung F, Lehnert H, Sprink T, Kohl C, Keilwagen J, Wilhelm R. 2020. Which Factors Affect the Occurrence of Off-Target Effects Caused by the Use of CRISPR/Cas: A Systematic Review in Plants. Frontiers in Plant Science 11.

Monroe JG, Srikant T, Carbonell-Bejerano P, Becker C, Lensink M, Exposito-Alonso M, Klein M, Hildebrandt J, Neumann M, Kliebenstein D, et al. 2022. Mutation bias reflects natural selection in Arabidopsis thaliana. Nature 602: 101–105.

Noël L, Moores TL, van der Biezen EA, Parniske M, Daniels MJ, Parker JE, Jones JDG. 1999. Pronounced Intraspecific Haplotype Divergence at the RPP5 Complex Disease Resistance Locus of Arabidopsis. The Plant Cell 11: 2099–2111.

Ordon J, Gantner J, Kemna J, Schwalgun L, Reschke M, Streubel J, Boch J, Stuttmann J. 2017. Generation of chromosomal deletions in dicotyledonous plants employing a user-friendly genome editing toolkit. The Plant Journal: For Cell and Molecular Biology 89: 155–168.

Ordon J, Kiel N, Becker D, Kretschmer C, Schulze-Lefert P, Stuttmann J. 2023. Targeted gene deletion with SpCas9 and multiple guide RNAs in Arabidopsis thaliana: four are better than two. Plant Methods 19: 30.

Ordon J, Martin P, Erickson JL, Ferik F, Balcke G, Bonas U, Stuttmann J. 2021. Disentangling cause and consequence: genetic dissection of the DANGEROUS MIX2 risk locus, and activation of the DM2h NLR in autoimmunity. The Plant Journal 106: 1008–1023.

Ossowski S, Schneeberger K, Lucas-Lledó JI, Warthmann N, Clark RM, Shaw RG, Weigel D, Lynch M. 2010. The Rate and Molecular Spectrum of Spontaneous Mutations in Arabidopsis thaliana. *Science (New York*, N.Y.) 327: 10.1126/science.1180677.

Park SH, Cao M, Pan Y, Davis TH, Saxena L, Deshmukh H, Fu Y, Treangen T, Sheehan VA, Bao G. 2022. Comprehensive analysis and accurate quantification of unintended large gene modifications induced by CRISPR-Cas9 gene editing. Science Advances 8: eabo7676.

Rabanal FA, Gräff M, Lanz C, Fritschi K, Llaca V, Lang M, Carbonell-Bejerano P, Henderson I, Weigel D. 2022. Pushing the limits of HiFi assemblies reveals centromere diversity between two Arabidopsis thaliana genomes. Nucleic Acids Research 50: 12309–12327.

Shen H, Strunks GD, Klemann BJPM, Hooykaas PJJ, de Pater S. 2017. CRISPR/Cas9-Induced Double-Strand Break Repair in Arabidopsis Nonhomologous End-Joining Mutants. G3 Genes|Genomes|Genetics 7: 193–202.

Shimada TL, Shimada T, Hara-Nishimura I. 2010. A rapid and non-destructive screenable marker, FAST, for identifying transformed seeds of Arabidopsis thaliana. The Plant Journal 61: 519–528.

Sinapidou E, Williams K, Nott L, Bahkt S, Tör M, Crute I, Bittner-Eddy P, Beynon J. 2004. Two TIR:NB:LRR genes are required to specify resistance to Peronospora parasitica isolate Cala2 in Arabidopsis. The Plant Journal 38: 898–909.

Smolka M, Paulin LF, Grochowski CM, Horner DW, Mahmoud M, Behera S, Kalef-Ezra E, Gandhi M, Hong K, Pehlivan D, et al. 2024. Detection of mosaic and population-level structural variants with Sniffles2. Nature Biotechnology 42: 1571–1580.

Sternberg SH, Redding S, Jinek M, Greene EC, Doudna JA. 2014. DNA interrogation by the CRISPR RNA-guided endonuclease Cas9. Nature 507: 62–67.

Sturme MHJ, van der Berg JP, Bouwman LMS, De Schrijver A, de Maagd RA, Kleter GA, Battaglia-de Wilde E. 2022. Occurrence and Nature of Off-Target Modifications by CRISPR-Cas Genome Editing in Plants. ACS Agricultural Science & Technology 2: 192–201.

Stuttmann J, Barthel K, Martin P, Ordon J, Erickson JL, Herr R, Ferik F, Kretschmer C, Berner T, Keilwagen J, et al. 2021. Highly efficient multiplex editing: one-shot generation of 8× Nicotiana benthamiana and 12× Arabidopsis mutants. The Plant Journal 106: 8–22.

Stuttmann J, Peine N, Garcia AV, Wagner C, Choudhury SR, Wang Y, James GV, Griebel T, Alcázar R, Tsuda K, et al. 2016. Arabidopsis thaliana DM2h (R8) within the Landsberg RPP1-like Resistance Locus Underlies Three Different Cases of EDS1-Conditioned Autoimmunity. PLOS Genetics 12: e1005990.

Sundaresan R, Parameshwaran HP, Yogesha SD, Keilbarth MW, Rajan R. 2017. RNA-Independent DNA Cleavage Activities of Cas9 and Cas12a. Cell Reports 21: 3728–3739.

Tsai SQ, Nguyen NT, Malagon-Lopez J, Topkar VV, Aryee MJ, Joung JK. 2017. CIRCLE-seq: a highly sensitive in vitro screen for genome-wide CRISPR–Cas9 nuclease off-targets. Nature Methods 14: 607–614.

Van de Weyer A-L, Monteiro F, Furzer OJ, Nishimura MT, Cevik V, Witek K, Jones JDG, Dangl JL, Weigel D, Bemm F. 2019. A Species-Wide Inventory of NLR Genes and Alleles in *Arabidopsis thaliana*. Cell 178: 1260–1272.e14.

Van Der Biezen EA, Freddie CT, Kahn K, Parker¶ JE, Jones JDG. 2002. Arabidopsis RPP4 is a member of the RPP5 multigene family of TIR-NB-LRR genes and confers downy mildew resistance through multiple signalling components. The Plant Journal 29: 439–451.

Vu GTH, Cao HX, Fauser F, Reiss B, Puchta H, Schubert I. 2017. Endogenous sequence patterns predispose the repair modes of CRISPR/Cas9-induced DNA double-stranded breaks in Arabidopsis thaliana. The Plant Journal 92: 57–67.

Zhang Y, Ge X, Yang F, Zhang L, Zheng J, Tan X, Jin Z-B, Qu J, Gu F. 2014. Comparison of non-canonical PAMs for CRISPR/Cas9-mediated DNA cleavage in human cells. Scientific Reports 4: 5405.

Zhou H, Liu B, Weeks DP, Spalding MH, Yang B. 2014. Large chromosomal deletions and heritable small genetic changes induced by CRISPR/Cas9 in rice. Nucleic Acids Research 42: 10903–10914.

