## Appendix 1 for "Analysis of *Sp*Cas9 on- and off-target effects in high efficiency multiplex editing in Arabidopsis"

**Appendix 1: Analysis of polymorphisms detected in *Δ13nlr* in a Col-0 individual**

Polymorphisms differentiating *Δ13nlr* from sequenced Col-0 genomes, as extracted from Table S1:

| chromosome | position | reference | alternative | length | length (alt) |
|---|---|---|---|---|---|
| Chr1 | 888694 | C | T | 1 | 1 |
| Chr1 | 4362700 | T | A | 1 | 1 |
| Chr1 | 6538500 | C | T | 1 | 1 |
| Chr1 | 7840015 | A | G | 1 | 1 |
| Chr1 | 8680485 | C | T | 1 | 1 |
| Chr1 | 8927804 | C | T | 1 | 1 |
| Chr1 | 11212342 | C | T | 1 | 1 |
| Chr1 | 11817700 | C | T | 1 | 1 |
| Chr1 | 12513589 | G | C | 1 | 1 |
| Chr1 | 12592042 | C | T | 1 | 1 |
| Chr1 | 12991433 | C | G | 1 | 1 |
| Chr1 | 16087780 | G | A | 1 | 1 |
| Chr1 | 16499211 | A | T | 1 | 1 |
| Chr1 | 16810895 | G | A | 1 | 1 |
| Chr1 | 17624941 | CAG | TAG | 3 | 3 |
| Chr1 | 20205381 | G | A | 1 | 1 |
| Chr1 | 20704510 | C | T | 1 | 1 |
| Chr1 | 23215331 | C | A | 1 | 1 |
| Chr1 | 25573615 | T | A | 1 | 1 |
| Chr2 | 713671 | G | A | 1 | 1 |
| Chr2 | 1057325 | A | G | 1 | 1 |
| Chr2 | 1310494 | C | T | 1 | 1 |
| Chr2 | 1618023 | T | C | 1 | 1 |
| Chr2 | 1856630 | C | T | 1 | 1 |
| Chr2 | 3857476 | AACG | AACA | 4 | 4 |
| Chr2 | 4932966 | G | A | 1 | 1 |
| Chr2 | 6923598 | A | T | 1 | 1 |
| Chr2 | 8551310 | A | G | 1 | 1 |
| Chr2 | 9565247 | A | T | 1 | 1 |
| Chr2 | 11434437 | A | C | 1 | 1 |
| Chr2 | 12286078 | T | C | 1 | 1 |
| Chr2 | 15240991 | T | A | 1 | 1 |
| Chr2 | 17314697 | G | A | 1 | 1 |
| Chr3 | 2261017 | C | T | 1 | 1 |
| Chr3* | 7580029 | C | T | 1 | 1 |
| Chr3 | 10713769 | C | T | 1 | 1 |
| Chr3 | 11597854 | G | A | 1 | 1 |
| Chr3 | 12718808 | G | A | 1 | 1 |
| Chr3 | 13300375 | CGTTC | TGTTC | 5 | 5 |

| | | | | | |
|---|---|---|---|---|---|
| Chr3 | 16934379 | C | T | 1 | 1 |
| Chr4 | 135769 | G | A | 1 | 1 |
| Chr4 | 3372325 | C | T | 1 | 1 |
| Chr4 | 4772360 | T | A | 1 | 1 |
| Chr4 | 10400586 | T | C | 1 | 1 |
| Chr4 | 10604994 | G | C | 1 | 1 |
| Chr4* | 13468905 | G | C | 1 | 1 |
| Chr4 | 16428223 | A | G | 1 | 1 |
| Chr5 | 355471 | A | G | 1 | 1 |
| Chr5 | 796263 | GTAGAATA | AAATGAGT | 8 | 8 |
| Chr5 | 796299 | G | A | 1 | 1 |
| Chr5 | 1848991 | T | C | 1 | 1 |
| Chr5 | 6349093 | C | T | 1 | 1 |
| Chr5 | 8227668 | C | G | 1 | 1 |
| Chr5 | 10913045 | A | T | 1 | 1 |
| Chr5 | 11818794 | T | C | 1 | 1 |
| Chr5 | 15716080 | C | T | 1 | 1 |
| Chr5 | 20686207 | G | C | 1 | 1 |
| Chr5 | 22441020 | C | A | 1 | 1 |
| Chr5 | 25742156 | G | A | 1 | 1 |
| Chr5 | 25810983 | T | G | 1 | 1 |

Color code:

Polymorphism detected in *Δ13nlr*, and also the Col-0 plant (derived from a seed batch different from the sequenced Col-2017/2019 control line) analyzed here.

Polymorphism detected in *Δ13nlr*, but not in later *Δnlr* lines. Accordingly, it segregated in the original wild type seed population used for transformation/transgenesis.

Polymorphism detected in *Δ13nlr*, but not the Col-0 plant analyzed here.

### Analysis of polymorphisms detected in *Δ13nlr* in a Col-0 individual

Amplicons spanning selected polymorphisms were Sanger-sequenced, and aligned to TAIR10 as reference. The position of the polymorphism as well as the oligonucleotides used for PCR amplification are indicated.

**Chr1:4362700; GACATCAAACGGGTCTAGGT, GGATGTTGTGAAATGCTGCA**

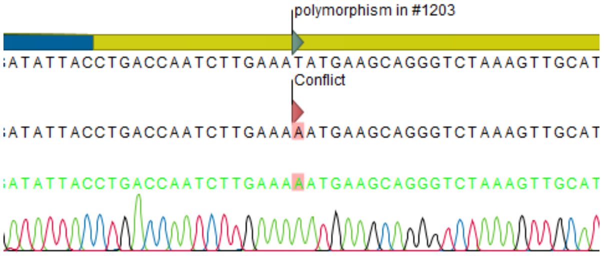


**Chr1:12592042; CGGTTGTTCGTAGTATGAATCAC, CGTCTTCACTCCGCTACTTC**

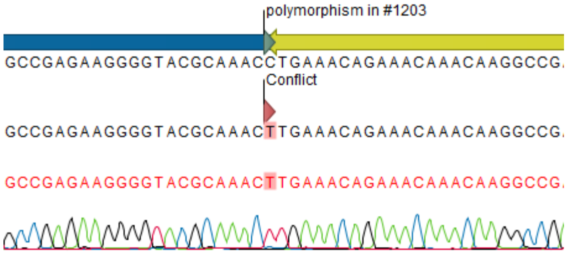

**Chr2:8551310; CGGGTCTAATCTGGGAAACG, AAAACCTCCCATGCCGATAA**

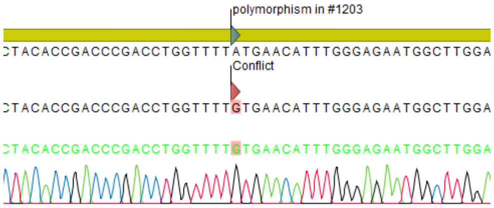


**Chr2:11434437; AGCTGCTCAACACTGTCAAA, GACATTCAAGTGGTCCCAAAC**

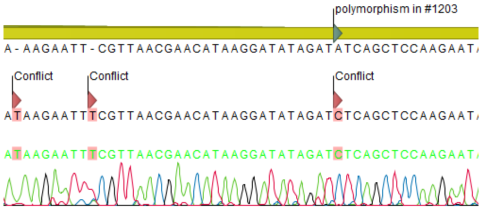


**Chr2:12286078; AGACTGAAGGCGAGAAAACG, AGAGCGCATAATCTTGTGACA**

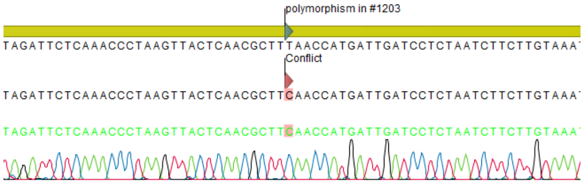


**Chr3: 2261017; TGGTTCAGAGGGCAATTTGG, TCGAAGAAGACTAGCCATAATCG**

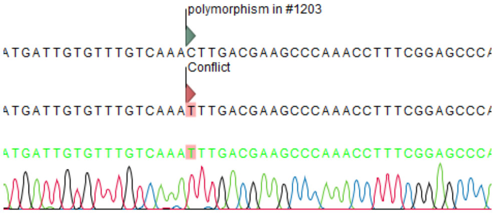


**Chr3:7580029; GAATGGAAGTTGGTCGTGCT, GCTGCATTTAGTTTTGGTTTTGA**

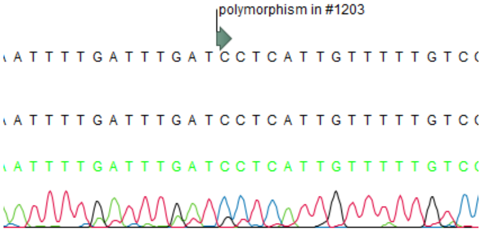


**Chr3:16934379: TGGTTAGTATTGGCGAATTGT, GTATCTCCTCATTGCAACATTCCG**

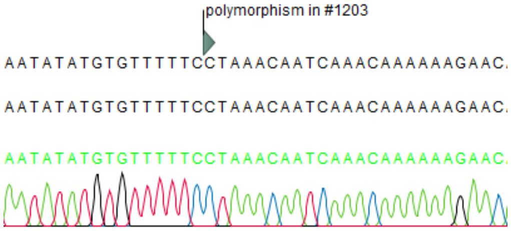


**Chr4:135769; GAAGGTTTTAGGAGCGTCTC, TCCGACAACCAACATCATCA**

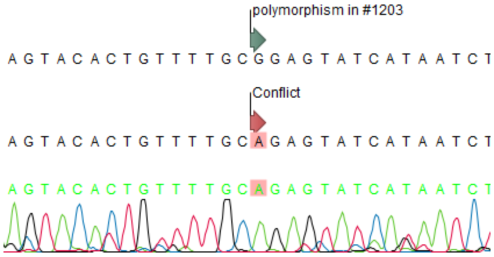


**Chr4:3372325; CCCATTGGACAATATGAAGTGCT, GGAACCCCACCAAGAGTATACT**

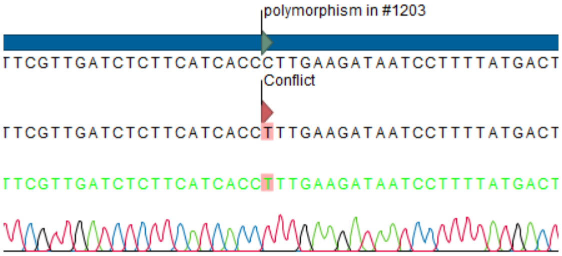


**Chr4:13468905; GCCCCTCCACCAATCTGTAT, CAGTGGCTAGCAAGTAGAGGA**

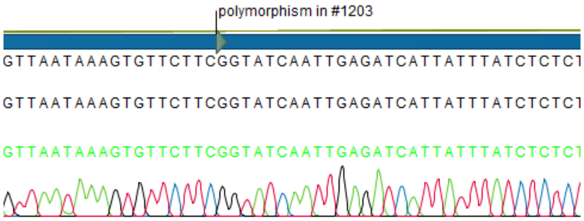


**Chr5:1848991; GGAGCCTCTTCTGCAATCAT, TTACTAGCCGCAAAACACGA**

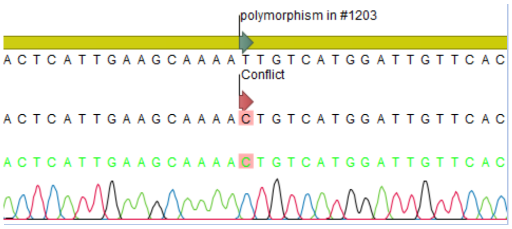


**Chr5:20686207; CTCCCAAGACCCAGGAAGAA, TCTCAACCACCTCCTGCTTT**

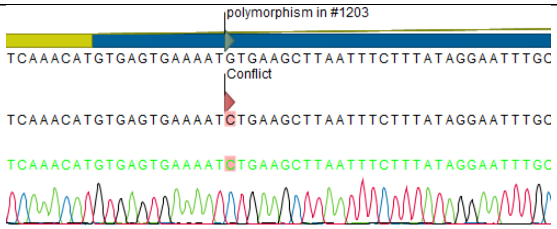
