## Supplementary material for "Analysis of *Sp*Cas9 on- and off-target effects in high efficiency multiplex editing in Arabidopsis": Figure S1

### Supplemental Figure S1

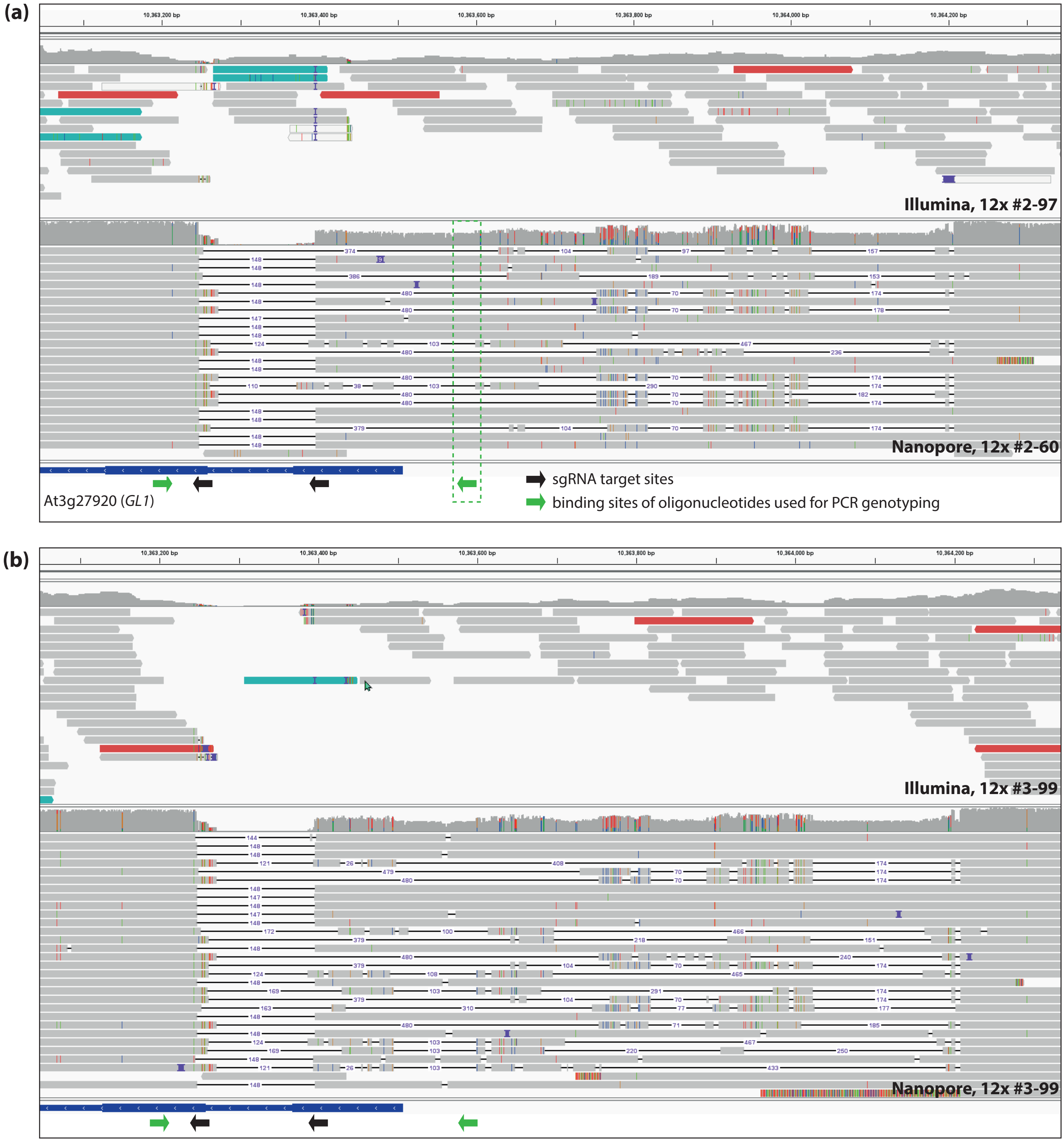


**Supplemental Figure S1: Read mappings at the *GL1* locus for 12x mutant lines.**

**a)** Illumina and Nanopore read mappings for the two indicated sister lines derived from 1681-2. The Nanopore mapping reveals that the respective individual was heterozygous for two different alleles: A deletion of 148 nt, and a complex allele summing up to a deletion of approximately 750 nt. The -750 nt deletion allele does not contain the binding site of a primer previously used for PCR-based genotyping, thus explaining lacking detection of this allele in the progenitor line 1681-2 during PCR genotyping (Stuttmann et al., 2021). The mapping of Illumina data, although it reveals that there are inconsistencies (e.g., blue shading indicates that two reads from a pair were not mapped in the expected orientations), does not allow correct determination of the genotype; none of the two possible alleles are suggested. No variant was detected at this locus by Freebayes with our parameters.
**b)** Screenshot of read mappings displayed in IGV as in a), but for line 1681-3-99. The Nanopore data allows the determination of the genotype of this individual at the *GL2* locus: As in 12x #2-60, a -148 deletion and a complex allele with deletion of approximately 750 nt. Illumina reads do not suggest any of these alleles; no variant was called by Freebayes.
