## Supplementary material for "Analysis of *Sp*Cas9 on- and off-target effects in high efficiency multiplex editing in Arabidopsis": Figure S2

### Supplemental Figure S2

(a)

(b)

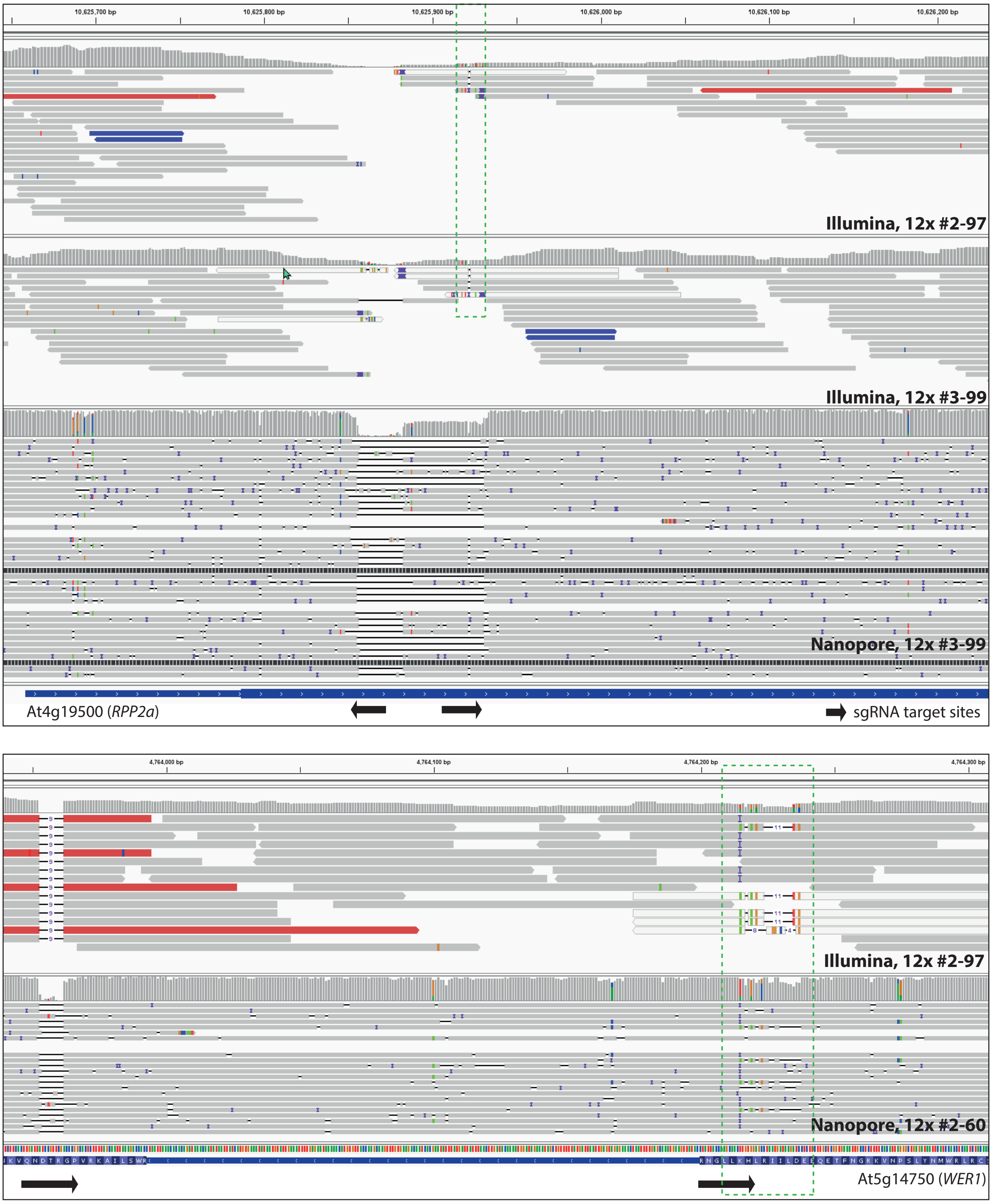


**Supplemental Figure S2: Read mappings at the *RPP2* and *WER1* loci for 12x mutant lines.**

**a)** Illumina and Nanopore read mappings for indicated lines at the *RPP2* locus. The Nanopore mapping (#3-99) reveals a -75 deletion and a second allele with -1 and -26 nt deletions. The same region is boxed (green) in the mappings of Illumina reads: Due to the large deletions, only very few reads were successfully mapped to the reference at this locus.

**b)** Read mappings displayed in IGV as in a), but at the *WER1* locus. A +1 insertion and a more complex allele (-8/-2/-1/-1, SNPs) is visible at target site 1 (green box) for line #2-60, sequenced by Nanopore technology. No variant was called at this locus.
