## Supplementary material for "Analysis of *Sp*Cas9 on- and off-target effects in high efficiency multiplex editing in Arabidopsis": Figure S3

### Supplemental Figure S3

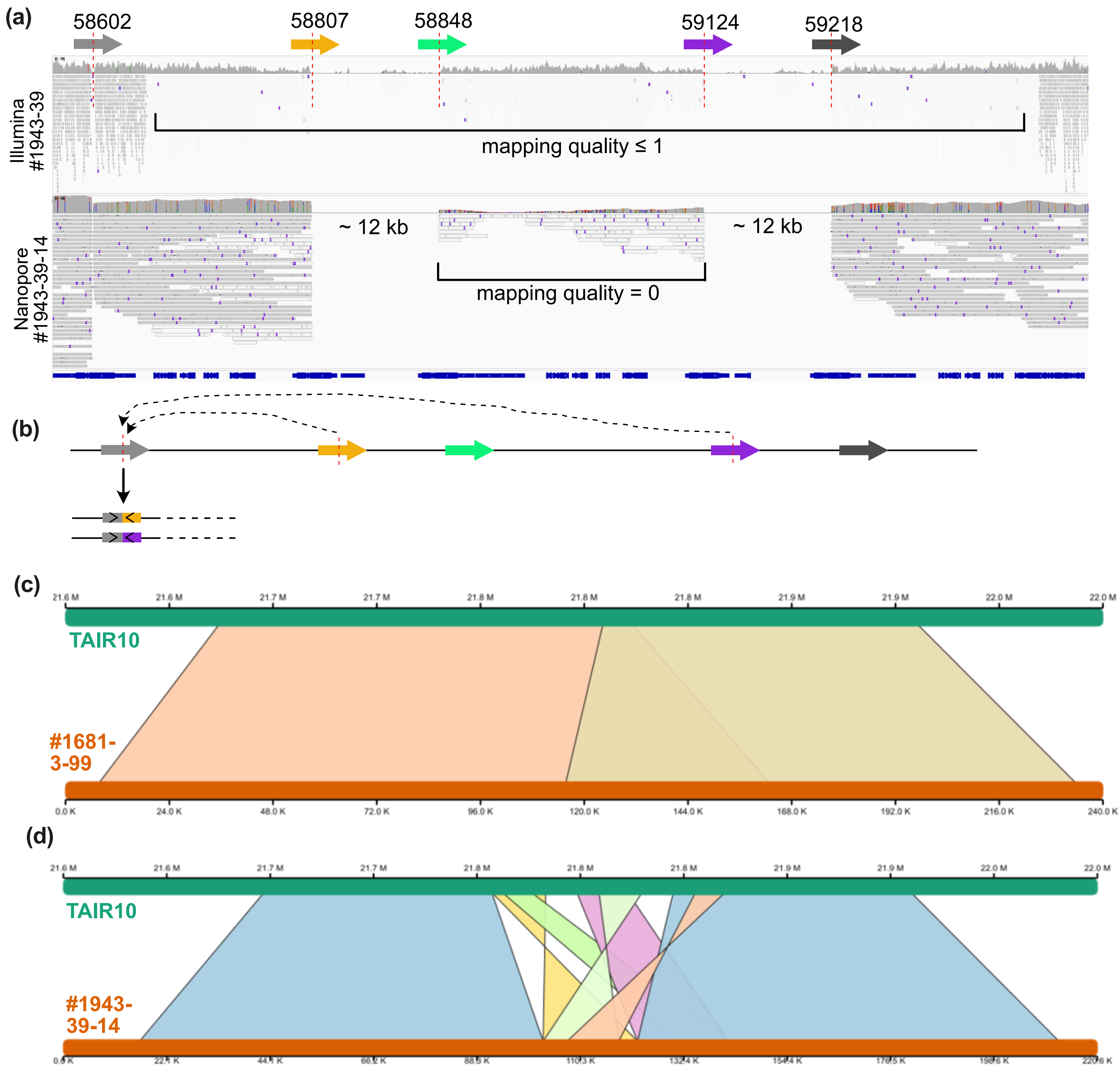


**Supplemental Figure S3: Analysis of recombination events at the *RPP7* locus.**

**a)** Mappings of Illumina and Nanopore reads for Δ*32nlr* lines visualized in IGV.
**b)** Possible inversion events suggested by analysis of Nanopore read ends at the breakpoint within At1g58602. Only the actual suggested event is shown, as the configuration of the remaining cluster remains unresolved.
**c)** Comparison of a *de novo* alignment of Nanopore reads from the *RPP7* region for a control line, #1681-3-99, without editing at the *RPP7* locus.
**d)** As in c), but reads from the Δ*32nlr* line #1943-39-14 were used for the *de novo* assembly.
