## Supplementary material for "Analysis of *Sp*Cas9 on- and off-target effects in high efficiency multiplex editing in Arabidopsis": Figure S4

### Supplemental Figure S4

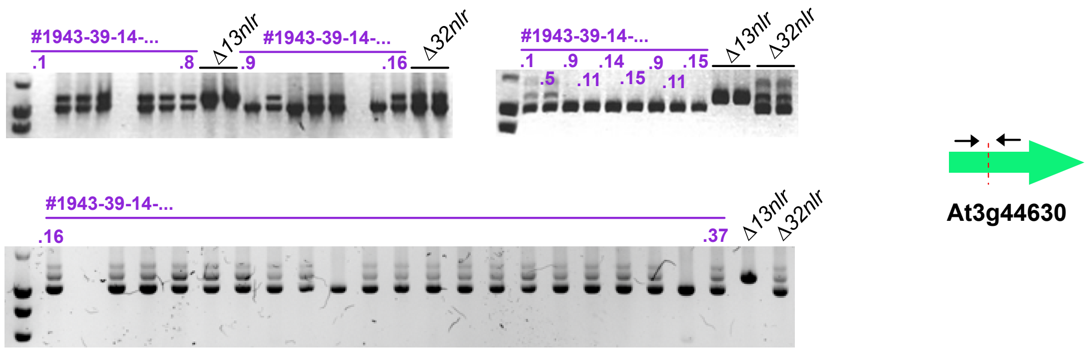


**Supplemental Figure S4: PCR-screening to identify plants homozygous for a single haplotype at the RPP1 locus.**

In total, 37 individuals derived from #1943-39-14 were analyzed using primer pair JS3099/3102, querying the CRISPR target site within At3g44630. DNA from individuals 14 and 15 was used for further analysis of junction sites.
