## Supplementary material for "Analysis of *Sp*Cas9 on- and off-target effects in high efficiency multiplex editing in Arabidopsis": Figure S5

### Supplemental Figure S5

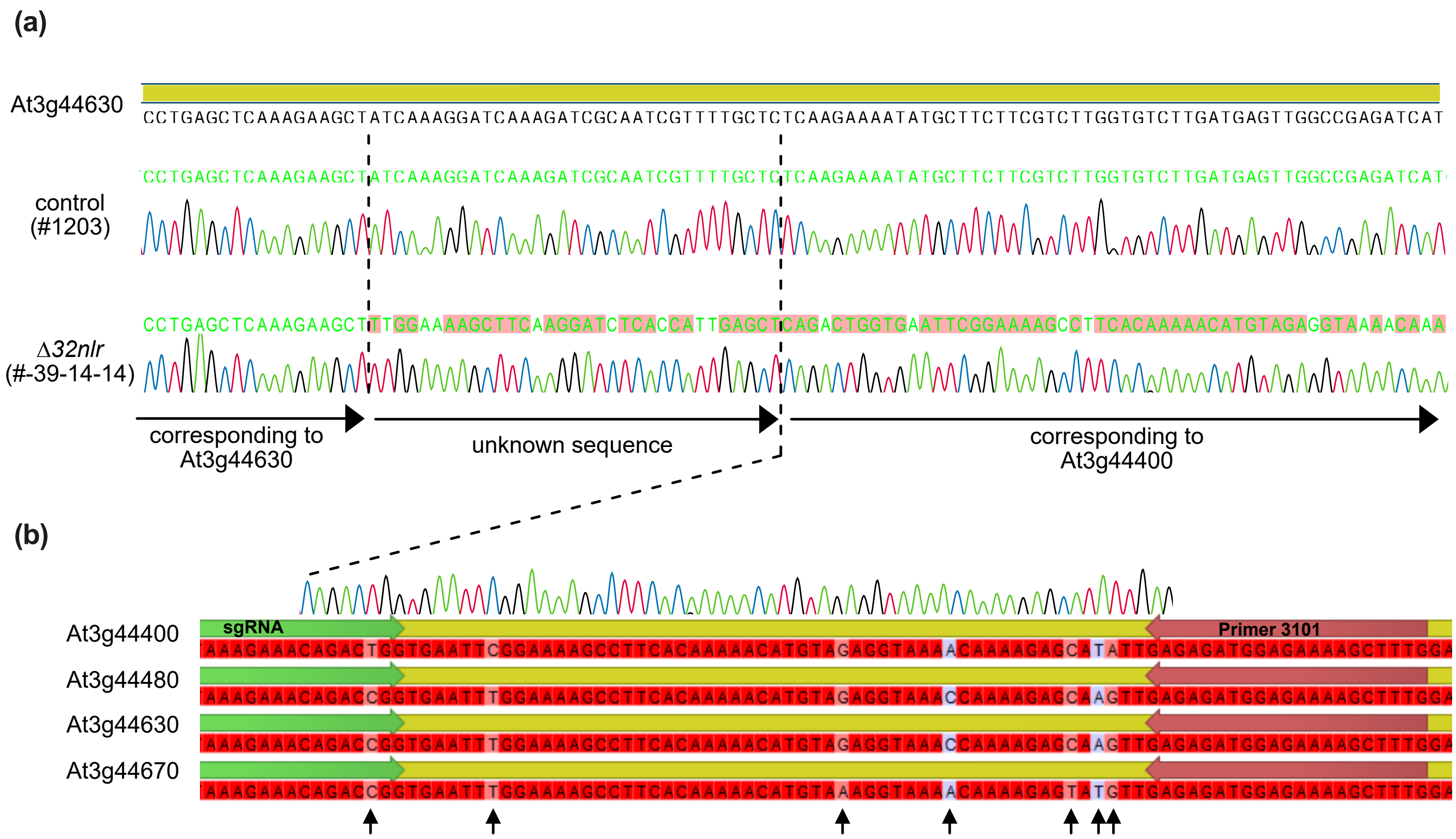


**Supplemental Figure S5: Inversion and fusion event involving At3g44630 and At3g44400 in Δ*32nlr*.**

**a)** Chromatograms from sequencing the PCR products obtained with oligonucleotides 3099/3102 from a control plant (#1203) and the Δ*32nlr* individual #1943-39-14-14, homozygous at the *RPP1* locus.
**b)** Alignment of a section of the chromatogram from the Δ*32nlr* individual from a) to an alignment of the *RPP1*-like genes. SNPs discriminating the genes are marked by arrows.
