## Supplementary material for "Analysis of *Sp*Cas9 on- and off-target effects in high efficiency multiplex editing in Arabidopsis": Figure S6

### Supplemental Figure S6

**(a)**

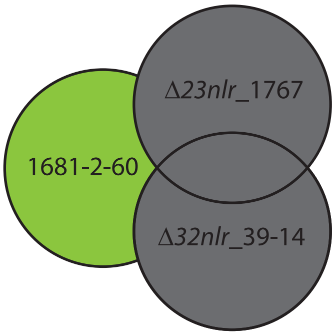


**1681-2-60** vs. Δ*23nlr*_1767 and Δ*32nlr*_39-14
**1681-3-99** vs. Δ*23nlr*_1767 and Δ*32nlr*_39-14
**Δ*23nlr*_1767** vs. 1681-2-60 and 1681-3-99
**Δ*32nlr*_39-14** vs. 1681-2-60 and 1681-3-99

**(b)**

1681-2-60

| Chr# | Position | Information | Format | Sample |
|---|---|---|---|---|
| Chr1 | 3886290 | SVTYPE=DEL;SVLEN=-10;END=3886300;SUPPORT=12;STRAND=+-;AF=0.444 | GT:GQ:DR:DV | 0/1:60:15:12 |
| Chr1 | 4556812 | SVTYPE=DEL;SVLEN=-13;END=4556825;SUPPORT=14;STRAND=+-;AF=0.560 | GT:GQ:DR:DV | 0/1:60:15:12 |
| Chr2 | 17946255 | SVTYPE=INS;SVLEN=43;END=17946255;SUPPORT=13;STRAND=+-;AF=1.000 | GT:GQ:DR:DV | 1/1:36:0:13 |

Δ*32nlr*_39-14

| Chr# | Position | Information | Format | Sample |
|---|---|---|---|---|
| Chr1 | 3886287 | SVTYPE=DEL;SVLEN=-12;END=3886299;SUPPORT=14;STRAND=+-;AF=0.389 | GT:GQ:DR:DV | 0/1:60:22:14 |
| Chr1 | 4556809 | SVTYPE=DEL;SVLEN=-12;END=4556821;SUPPORT=18;STRAND=+-;AF=0.600 | GT:GQ:DR:DV | 0/1:60:12:18 |
| Chr2 | 17946217 | SVTYPE=INS;SVLEN=46;END=17946217;SUPPORT=16;STRAND=+-;AF=1.000 | GT:GQ:DR:DV | 1/1:44:0:16 |

**(c)**

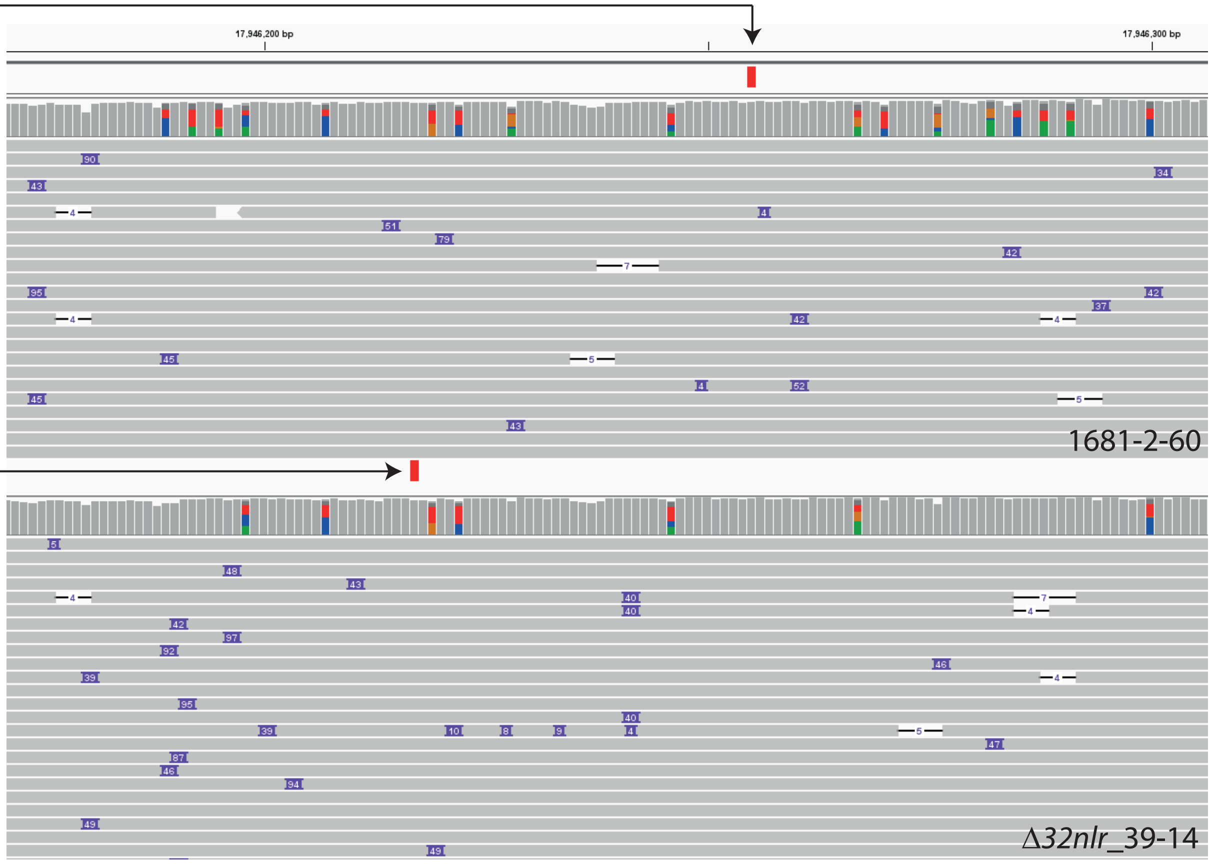


**Supplemental Figure S6: Intersection of variants detected by Sniffles2.**

**a)** Intersection of variants with those of genetically independent lines. One example shown as Venn diagram: Only variants unique to 1681-2-60 (green area) were retained.
**b)** Sniffles2 often assigned different coordinates to similar variants in datasets. Therefore, variants were excluded during intersection when a variant with coordinates +/-20 was detected in an independent lines. The first two variants for each dataset represent examples of variants that were excluded by this procedure. The last line shows a variant that was retained, but discarded after inspection in IGV. Similar variants are detecetd in both mappings, but coordinates differ by ~40 nt.
**c)** Read mappings showing the variant from **b)** that was excluded after inspection in IGV. Violet boxes on individual Nanopore reads indicate insertions, mostly around 40 nt. Sniffles2 called an insertion in both lines (red boxes above read coverage plot; variant call file displayed in IGV), but with different coordinates, as shown in **b)**. Arrows indicate correspondance between variants shown in **b)** as table and in **c)** together with the IGV-displayed read mappings.
